# PCNA monoubiquitination is regulated by diffusion of Rad6/Rad18 complexes along RPA filaments

**DOI:** 10.1101/2020.08.11.247064

**Authors:** Mingjie Li, Bhaswati Sengupta, Stephen J. Benkovic, Tae Hee Lee, Mark Hedglin

## Abstract

Translesion DNA synthesis (TLS) enables DNA replication through damaging modifications to template DNA and requires monoubiquitination of the PCNA sliding clamp by the Rad6/Rad18 complex. This posttranslational modification is critical to cell survival following exposure to DNA damaging agents and is tightly regulated to restrict TLS to damaged DNA. RPA, the major single strand DNA (ssDNA) binding protein, forms filaments on ssDNA exposed at TLS sites and plays critical yet undefined roles in regulating PCNA monoubiquitination. Here, we utilize kinetic assays and single molecule FRET microscopy to monitor PCNA monoubiquitination and Rad6/Rad18 complex dynamics on RPA filaments, respectively. Results reveal that a Rad6/Rad18 complex is recruited to an RPA filament via Rad18•RPA interactions and randomly translocates along the filament. These translocations promote productive interactions between the Rad6/Rad18 complex and the resident PCNA, significantly enhancing monoubiquitination. These results illuminate critical roles of RPA in the specificity and efficiency of PCNA monoubiquitination.

## INTRODUCTION

The B-family DNA polymerases (pols) ε and δ replicate the majority of the human genome and achieve optimal processivity by anchoring to PCNA sliding clamps encircling primer/template (P/T) junctions^1^. These “replicative” pols have very stringent polymerase domains and 3’ to 5’ exonuclease (“proofreading”) domains that collectively ensure accurate replication of native template bases during S-phase of the cell cycle. However, DNA is continuously damaged by covalent modifications from reactive metabolites and environmental mutagens, such as ultraviolet radiation (UVR), and the replicative pols cannot accommodate damaged template bases (i.e. lesions). Consequently, primer extension by these pols stalls upon encountering a lesion, leading to persistent exposure of the template strand downstream of the lesion. At UVR-induced lesions, the exposed templates range in length from 150 nt to 1250 nt, with the latter accounting for 65%^2,3^. RPA immediately coats the exposed templates (1 RPA/30 + 2 nt)^4–6^, forming elongated and persistent RPA filaments that protect the underlying ssDNA from degradation, prevents formation of alternative DNA structures^7^, and blocks diffusion of PCNA along the damaged template^8–10^. Failure to restart primer extension on damaged templates often results in double-strand breaks that may lead to gross chromosomal rearrangements (GCRs), cell-cycle arrest, and cell death. These stalling events may be overcome by translesion DNA synthesis (TLS) where specialized TLS pols bind the resident PCNA and extend the stalled primer across and beyond the DNA lesion, allowing DNA synthesis by a replicative pol to resume downstream of the lesion^7^. Characterized by a more “open” DNA polymerase active site and the lack of an associated proofreading activity, a single TLS pol can accommodate multiple DNA lesions, albeit with varying fidelities^11^. Hence, tight regulation is required to limit the frequency and extent of TLS.

In humans, TLS requires the covalent attachment of single ubiquitin moieties (i.e. monoubiquitination) to lysine residues K164 of PCNA sliding clamps encircling stalled P/T junctions^12^. This critical posttranslational modification (PTM) is fully conserved in eukaryotes and catalyzed by a complex comprised of Rad6 and Rad18 proteins. The former is an E2 ubiquitin conjugating enzyme that covalently attaches a ubiquitin to PCNA and the latter is a RING E3 ubiquitin ligase that delivers Rad6 to a PCNA target^13^. PCNA monoubiquitination contributes to cell survival following exposure to DNA-damaging agents such as UVR and, hence, must be tightly-regulated as dysfunction can selectively propagate cells with increased mutagenesis due to aberrant TLS^7,13^. Currently, it is unclear how the activity of the Rad6/Rad18 complex is regulated, particularly regarding the roles of RPA. Recent studies revealed that Rad6/Rad18 complexes directly interact (via Rad18) with RPA on ssDNA and this non-specific interaction is required for PCNA monoubiquitination^10,14,15^. A pressing issue that has remained unresolved is the functional role(s) of non-specific Rad18•RPA interactions in PCNA monoubiquitination. To investigate this, we utilized transient-state kinetic studies and single molecule FRET (smFRET) TIRF microscopy to directly monitor PCNA monoubiquitination and the dynamics of Rad6/Rad18 complexes on RPA filaments, respectively. Results from thorough experiments reveal that a Rad6/Rad18 complex is directly recruited to an RPA filament (via Rad18•RPA interactions) and then randomly translocates along the filament by one-dimensional, thermal-driven diffusion. These translocations promote productive interactions between the Rad6/Rad18 complex and the resident PCNA, significantly enhancing monoubiquitination. These results illuminate critical roles of RPA in the specificity and efficiency of PCNA monoubiquitination.

## RESULTS

### Monoubiquitination of PCNA encircling a P/T junction is promoted by the adjacent RPA filament

During a catalytic cycle (**Fig. 1A**), a Rad6/Rad18 complex charged with ubiquitin must first locate and engage its target substrate in a productive complex. The target substrate (S) is a PCNA encircling a P/T junction and is referred to herein as simply “loaded PCNA”. In the subsequent chemical step, the Rad6/Rad18 complex covalently attaches the associated ubiquitin to the engaged target, forming product. The apo Rad6/Rad18 complex devoid of ubiquitin then disengages from the product and turns over. To investigate the effect of non-specific Rad18•RPA interactions on catalysis, we monitored the transient-state kinetics of PCNA monoubiquitination on DNA that mimics stalled P/T junctions (**Fig. 1B**). The duplex regions are identical and the total lengths of the poly(dT) ssDNA regions are either 33 or 171 nt, which accommodates 1 and ~6 RPA molecules, respectively^4–6^. All experiments were carried out such that the concentrations of excess RPA are identical to account for any effects of “free” RPA in solution on PCNA monoubiquitination^10^. Charged Rad6/Rad18 is limiting, high concentrations of loaded PCNA are utilized, and only 5.53 + 1.05% of the reactions are monitored. Under these conditions, double binding events are very rare (< 0.15%), the pre-charged Rad6/Rad18 complexes may saturate with substrate (*k*_on_[S]) prior to the chemical step (*k*_chem_), and product release (*k*_release_) is irreversible (**Fig. 1A**). For all conditions analyzed (**Fig. 1B**), products accumulate during an initial burst of enzyme activity (i.e., burst phase) and then increase at a slower, constant rate (i.e., steady state phase) (**Fig. 1C**). This biphasic behavior indicates that all steps up to and including monoubiquitination of PCNA (*k*_on_|S| and *k*_chem_) are comparable to any subsequent steps during turnover (*k*_release_ and after). Fitting the steady state phases to linear regressions yields the amplitudes (A_0_) for the burst phases and the steady state rates (*v*_ss_) for turnover (reported in **Table 1**). The observed rate constants for PCNA monoubiquitination (*k*_mu,obs_) and turnover (*k*_cat_) are calculated from the values for *v*_ss_ and A_0_ (see **Methods**)^16^. *k*_cat_ reflects the release of product, re-charging of the apo Rad6/Rad18 complex with ubiquitin, or a combination of both steps (**Fig. 1A**). The products (monoubiquitinated PCNA) and apo Rad6/Rad18 complexes are identical for all conditions. Thus, it is expected that *k*_cat_ remains constant.

**Figure 1.**
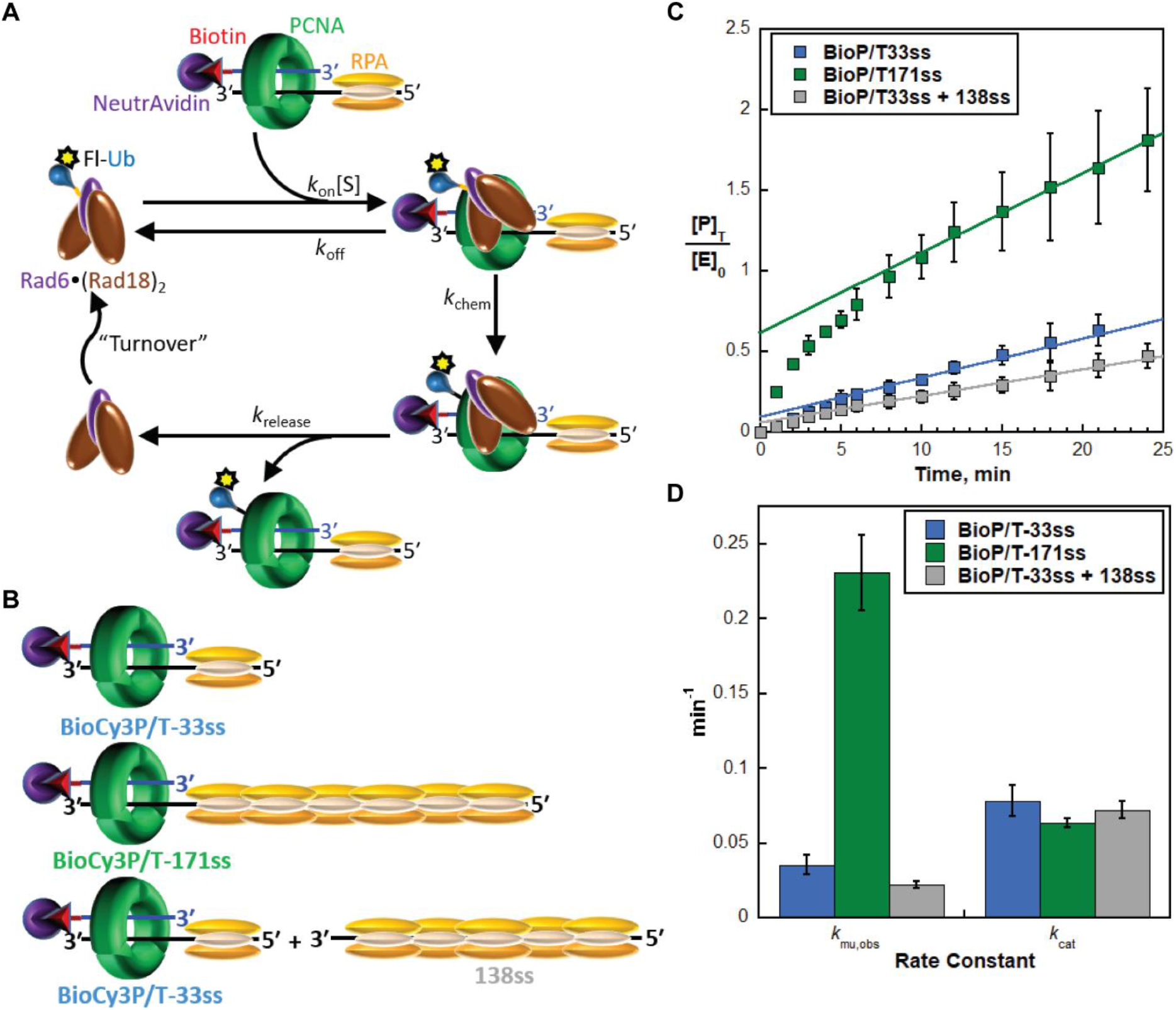
Transient-state kinetic analyses reveal a critical role for RPA in monoubiquitination of PCNA by the Rad6/Rad18 complex. (**A**) Schematic representation of the catalytic cycle of a charged Rad6/Rad18 complex and the experiments to monitor PCNA monoubiquitination. A PCNA is pre-assembled on a BioP/T DNA substrate in the presence of RPA. Both RPA and the biotin/neutravidin complexes serve to prevent to PCNA from sliding off the DNA. The Rad6/Rad18 complex charged with fluorescently-labeled ubiquitin is then added and monoubiquitination of target proteins is monitored over time. (**B**) Schematic representations of PCNA assembled onto the BioP/T DNA substrates. (**C**) Extents of PCNA monoubiquitination. The concentrations of monoubiquitinated PCNA clamps are divided by the initial concentration of charged Rad6/Rad18 and plotted as function of time. Data represent the average ± S.D. of three independent experiments. For each condition, the linear phase is fit to a linear regression that is extrapolated back to the y-axis. The y-intercept and the slope of the fit represent the amplitude (A_0_) and the steady state rate (*v*_ss_), respectively. (**D**) Kinetic analyses. Values for the rate constants *k*_mu,obs_ (■) and *k*_cat_ (◆) are calculated from the values for A_0_ and *v*_ss_ determined from panel **A** (and reported in **Table** 1) and plotted for each condition. *k*_mu,obs_ is dependent on the length of RPA molecules adjacent to the target PCNA. *k*_cat_ remains constant.

**Table 1.**
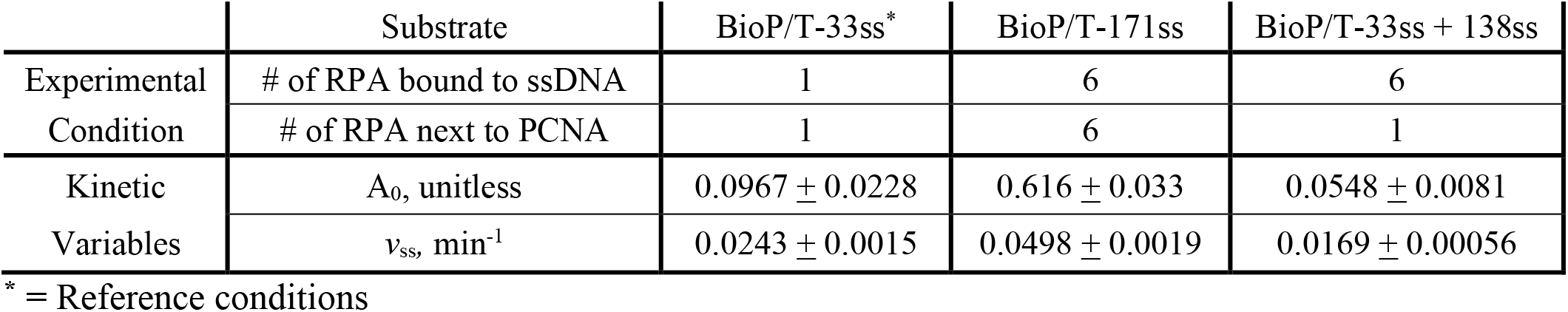
Kinetics obtained from transient state kinetic assays reveal a critical role for RPA in monoubiquitination of PCNA by the Rad6/Rad18 complex

A single RPA molecule resides next to a PCNA encircling the BioP/T-33ss DNA (**Fig. 1B**). Thus, the target (PCNA) and non-specific binding sites (RPA) are equally abundant on the P/T DNA and a Rad6/Rad18 complex has an equal probability of binding either from solution. For this reference condition *k*_mu,obs_ = 0.231 + 0.025 min^−1^ and *k*_cat_ = 0.0634 + 0.0029 min^−1^. On the BioP/T-171ss DNA (**Fig. 1B**), the number of RPA molecules residing next to loaded PCNA is increased to 6. Here, a Rad6/Rad18 complex is six times more likely to initially engage a non-specific site along the RPA filament rather than the loaded PCNA target. In other words, a Rad6/Rad18 complex is first localized/recruited to the RPA filament before engaging loaded PCNA and catalyzing monoubiquitination^14^. As observed in **Fig. 1D**, *k*_mu,obs_ is stimulated by a factor of 6.55 + 0.03 relative to the reference condition but *k*_cat_ is unaffected. The latter supports the validity of the approach. *k*_mu,obs_ is dependent on formation of the productive complex (*k*_on_[S]) and the chemical step (*k*_chem_) (**Fig. 1A**). The concentration of the target substrate ([S]) and *k*chem are identical for both DNAs as the number of RPA molecules residing next to the P/T junctions has no effect on the amount of loaded PCNA or its orientation on DNA (**Supplementary Fig. S3**)^10^. This suggests that *k*_mu,obs_ is rate-limited by *k*_on_ and, hence, overabundant non-specific sites (i.e., an RPA filament) adjacent to a loaded PCNA target stimulate *k*_mu,obs_ by increasing *k*_on_. A near stoichiometric increase in kmu,obs (6.55-fold) with the 6-fold overabundance of RPA molecules also supports this conclusion. To confirm this further, we repeated the experiments by separating the RPA filament from the P/T DNA (BioP/T-33ss + 138ss, **Fig. 1B**). Here, the loaded PCNA (on the BioP/T-33ss) and the detached RPA filament (on the 138ss) are stoichiometric. Hence, the number of RPA molecules bound to ssDNA is increased from one to six compared to the reference condition but still only a single RPA molecule resides next to a PCNA. As observed in **Fig. 1D**, the significant stimulation of *k*_mu,obs_ on the BioP/T-171ss DNA disappears when the RPA filament is not physically connected to the P/T junction encircled by PCNA. Again, *k*_cat_ remains constant, confirming the validity of the experimental approach (**Figure 1D**). Altogether, the results presented thus far indicate that recruitment of a Rad6/Rad18 complex to an RPA filament adjacent to loaded PCNA stimulates monoubiquitination (*k*_mu,obs_) by promoting formation of a productive complex (*k*_on_).

### Rad6/Rad18 complexes diffuse randomly along RPA filaments

In assays carried out with the BioP/T-171ss DNA, a Rad6/Rad18 complex in solution most likely engages a non-specific site along the RPA filament that is separated from the loaded PCNA target by one or more intervening RPA molecules. This recruitment significantly enhances PCNA monoubiquitination (*k*_mu,obs_) by promoting formation of the productive complex (*k*_on_) (**Fig. 1**). For catalysis to occur after recruitment, the engaged Rad6/Rad18 complex must transfer from a distal, non-specific site along the RPA filament to the PCNA target and the RPA filament must provide a pathway for transfer. A possible mechanism is direct transfer via ssDNA looping. However, RPA filaments linearize the underlying ssDNA in a rigid rod type structure by engaging the ssDNA in an elongated manner that extends the bound sequence and increases its bending rigidity 2 – 3 fold^17,18^. Alternatively, a Rad6/Rad18 complex may translocate along the RPA filament by random, thermal-driven diffusion. In other words, a Rad6/Rad18 complex diffuses towards and away from loaded PCNA during each engagement with the adjacent RPA filament. Such movements promote formation of the productive complex (*k*_on_) by decreasing the time to locate a PCNA target and/or permitting multiple encounters with a loaded PCNA target during each interaction with a DNA. To directly observe diffusion of Rad6/Rad18 complexes, we investigated the dynamics of Rad6/Rad18 complexes on RPA filaments by smFRET TIRF microscopy (**Fig. 2A**). The P/T DNA (BioCy3P/T-70ss, **Supplementary Fig. 1**) contains a biotin tag at the blunt duplex end, a Cy3 dye at the P/T junction, and accommodates two RPA molecules on the ssDNA^4–6^. The ssDNA is saturated with RPA and extended into an elongated, rigid filament^19^. Rad6 was first labeled with a single, N-terminal Cy5 dye (**Supplementary Fig. 4**) and then re-constituted with Rad18 to form the Rad6/Rad18 complex^20^. In this setup, smFRET occurs when Cy5-Rad6 (FRET acceptor) is in close proximity to the Cy3 dye (FRET donor) at the P/T junction. smFRET is only observed (i.e., t_on_) when both Rad18 and RPA are included (**Fig. 2B**); smFRET events are not detected during ~5520 minutes of total observation when either Rad18 or RPA are omitted. Altogether, this confirms that; 1) Rad18 functionally interacts with Cy5-Rad6 in a manner that directs Cy5-Rad6 to the vicinity of the P/T junction^13^ and; 2) the Rad6/Rad18 complex has exceptionally weak affinity for naked ssDNA and, hence, is directly recruited to DNA by RPA filaments^10,14^. Next, we analyzed the smFRET trajectories.

**Figure 2.**
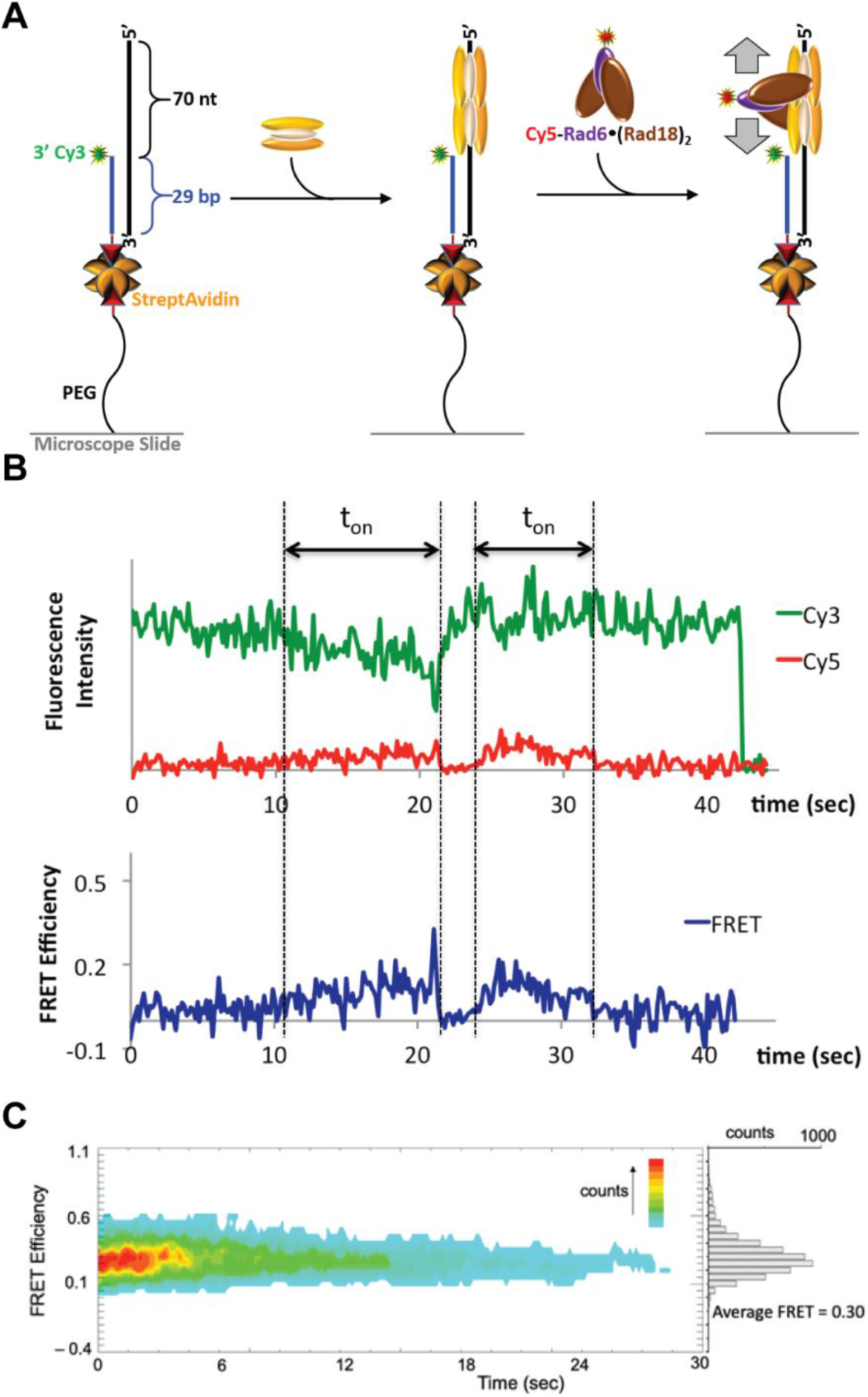
smFRET analyses reveal random diffusion of Rad6/Rad18 complexes on RPA filaments. (**A**) Experimental schematic to monitor translocation of a Rad6/Rad18 complex along RPA filaments via smFRET. The BioCy3P/T-70ss DNA substrate is immobilized on a microscope slide surface via biotin/streptavidin conjugation and the ssDNA region is saturated with two RPA molecules. Cy5-Rad6/Rad18 is injected and smFRET is monitored over time. (**B**) An example of a time trajectory shows the fluctuating smFRET efficiency during τ_on_, indicating that a Rad6/Rad18 complex translocates along an RPA filament (right). (**C**) Collective smFRET efficiency time trajectories (n = 88) synchronized at the starting point of smFRET events (i.e. starting points of the τon windows shown in panel **B**) were overlaid (*Left*). The average smFRET efficiency (indicated) is calculated from the histogram of the smFRET efficiencies observed during the τ_on_ windows (*Right*).

As depicted in the sample smFRET trajectory (**Fig. 2B**), the efficiencies during t_on_ rapidly fluctuate with time, lacking defined, stable conformational states. Such behavior is consistent with the translocation of a Cy5-Rad6/Rad18 complex towards and away from the Cy3-P/T junction via diffusion along the RPA filament. Alternative explanations are that a Cy5-Rad6/Rad18 complex is stably bound to a position on the RPA filament and the fluctuations arise due to the conformational dynamics of either the P/T junction or the engaged RPA molecules that enable contact between the FRET dyes. However, the conformational dynamics of DNA junctions are very fast and average out during the measurements with our signal integration time of 150 ms^21,22^. Also, the average dwell times (t_on_) of RPA DNA-binding domains (DBD) that undergo microscopic dissociation/re-association on ssDNA range from 300 ms – 1 s and such events would be clearly visible with the time resolution of the current experiments^23^. Although it cannot be ruled out that Rad6/Rad18 complexes affect these microscopic dissociation/re-association events, defined conformational states are not observed in the smFRET efficiencies (**Fig. 2B**). Altogether, this suggests that the fluctuations of smFRET efficiencies during t_on_ reflect the translocation of a Rad6/Rad18 complex towards and away from the P/T junction via diffusion along the RPA filament. Next, we investigated the directionality and speed of Rad6/Rad18 diffusion.

The 88 detectable smFRET events from the collective time trajectories were synchronized at the starting points of the t_on_ windows and overlaid (**Fig. 2C**, left). Histograms of the smFRET efficiencies observed during the t_on_ windows were constructed to show the distribution of the FRET efficiency (**Fig. 2C**, right). The average FRET efficiency calculated from the distribution is 0.30 and the FRET efficiency fluctuates randomly about this value with no sign of discrete FRET states. Together, this indicates that Rad6/Rad18 complexes diffuse randomly along the RPA filament, encountering the P/T junction multiple times during each binding interaction, in accordance with the proposed model. The changes in FRET efficiencies between two points separated by 3.45 ~ 5.4 sec (n = 55~174) were utilized to calculate the mean squared displacements (MSD) using a F örster radius of 5.4 nm for Cy3/Cy5^24^. Shorter t_on_ windows were removed from the analysis because the FRET dynamics within a short time window are dominated by noise. Longer t_on_ windows were also removed from the analysis because the sample size with longer time windows becomes too small. We found the analysis window resulting in the highest R^2^ value of the fitting. MSDs were plotted as a function of diffusion time (t) and fit to a linear regression, MSD = 2Dt, where D represents the 1D diffusion coefficient. This yields a low estimate for a 1D diffusion coefficient of 0.11 + 0.004 nm^2^s^−1^ (R^2^ = 0.80).

Collectively, the results presented in **Fig. 2** confirm that Rad6/Rad18 complexes are directly recruited to the vicinity of P/T junctions by the adjacent RPA filaments^10,14,15^ and reveal that Rad6/Rad18 complexes randomly diffuse along RPA filaments. These behaviors collectively enhance the catalytic activity of Rad6/Rad18 complexes by promoting formation of productive complexes (*k*_on_) with PCNA encircling P/T junctions (**Fig. 1**). These unforeseen results illuminate the undefined roles of Rad18 oRPA interactions in regulating PCNA monoubiquitination, as discussed below.

## Discussion

Recent studies revealed that Rad6/Rad18 complexes directly interact (via Rad18) with RPA on ssDNA and these non-specific interactions are required for PCNA monoubiquitination^10,14,15^. A pressing issue that has remained unresolved is the functional role(s) of non-specific Rad18•RPA interactions in PCNA monoubiquitination. In the present study, we utilized transient-state kinetic studies and single molecule FRET (smFRET) TIRF microscopy to directly monitor PCNA monoubiquitination and RPAoRad18 interactions on ssDNA, respectively. Results from thorough experiments revealed that; 1) Rad6/Rad18 complexes translocate along RPA filaments by random, thermal-driven diffusion, and; 2) these translocations significantly enhance monoubiquitination of PCNA encircling distal P/T junctions. These results reveal a catalytic mechanism that is unique to the Rad6/Rad18 complex among PCNA-modifying enzymes and, to our knowledge, the first example of ATP-independent translocation of a protein complex along a protein filament. Furthermore, this unique mechanism accounts for the many challenges that arise *in vivo,* namely selectivity/specificity and efficiency.

Monoubiquitination of PCNA elicits DNA synthesis by error-prone TLS pols and, hence, must be restricted to PCNA encircling P/T junctions stalled at DNA lesions, such as those generated by UVR exposure. UVR fluences similar to what an individual experiences from one hour of mid-day sun generate 1.6 to 2.2 million lesions in a human cell, with the vast majority (67 – 83%) undergoing very slow repair and persisting into S-phase ^25,26^. However, DNA replication in a human cell emanates from 13 to 22 million P/T junctions, each encircled by PCNA. Thus, very few (< 10 %) loaded PCNA clamps will ever encounter and subsequently idle at a UVR-induced lesion under physiologically-relevant conditions^25,27,28^. So, how is Rad6/Rad18 activity restricted to a such a small minority of loaded PCNA clamps? Compounding this issue is the relative abundance of Rad6/Rad18 complexes. ~50 proteins interact with loaded PCNA during S-phase in human cells and many are substantially enriched (as high as ~80-fold) at P/T junctions stalled at UVR-induced lesions^29^. However, the abundance of Rad6/Rad18 complexes is maintained at a low level (< ~795 /cell) and does not change following UVR exposure^30,31^. How can Rad6/Rad18 complexes effectively compete with the vast overabundance of competitive PCNA-binding proteins in human cells^32^? The present study along with previous work from our group and others suggests that selectivity and efficiency of PCNA monoubiquitination is achieved through non-specific Rad18•RPA interactions. On native DNA templates, RPA filaments adjacent to progressing P/T junctions are short and transient^33^ due to the minimal exposure of native ssDNA templates^7^ and their rapid conversion to double-stranded DNA duplexes by the replicative pols ε and δ (**Fig. 3**, left)^34^. Here, Rad18•RPA interactions on ssDNA are prohibited and Rad6/Rad18 complexes must engage loaded PCNA directly from solution. These events are inhibited *in vivo*^35^, likely by the continuous, rapid movement of PCNA engaged with replicating DNA polymerases^10^ and the vast overabundance of competitive PCNA-binding proteins in human cells^29–32^. In support of this, Rad18 is diffusely distributed throughout the nucleus during S-phase in mock UVR-treated human cells^35^ and overexpression of Rad18 is required for PCNA monoubiquitination under these conditions^14^. In stark contrast, P/T junctions stalled at UVR-induced lesions generate RPA filaments that range in length from 5 - 42 RPA molecules^2–6^ and persist for > 8h^36^. The unique properties of these RPA filaments promote Rad18 oRPA interactions and selectively localize Rad6/Rad18 complexes to rare target sites independently of PCNA binding (**Fig. 3**, right). This avoids a biased competition with most cellular proteins that must localize via direct binding to loaded PCNA ^14^. Given the relative low abundance of Rad6/Rad18 complexes^30,31^ and the extended lengths of RPA filaments generated at UVR-induced lesions^2–6^, a single Rad6/Rad18 complex is initially recruited to a random position along an RPA filament that is distal to the loaded PCNA target. Once engaged, the Rad6/Rad18 complex translocates randomly along the RPA filament by thermal-driven diffusion. In other words, the non-specific binding interactions with RPA molecules are correlated, allowing a Rad6/Rad18 complex to engage many RPA molecules during each encounter with an RPA filament. A correlated search of non-specific sites for a target site is more efficient than a non-correlated search^37^. Furthermore, as the resident PCNA is being stochastically sampled by the nucleoplasmic pool of PCNA-binding proteins, diffusion of the Rad6/Rad18 complex along the adjacent RPA filament selectively elevates the relative frequency of collisions between the loaded PCNA target and the Rad6/Rad18 complex. Together, this promotes monoubiquitination of PCNA encircling stalled P/T junction despite the relatively low abundance of Rad6/Rad18 complexes^30,31^.

**Figure 3.**
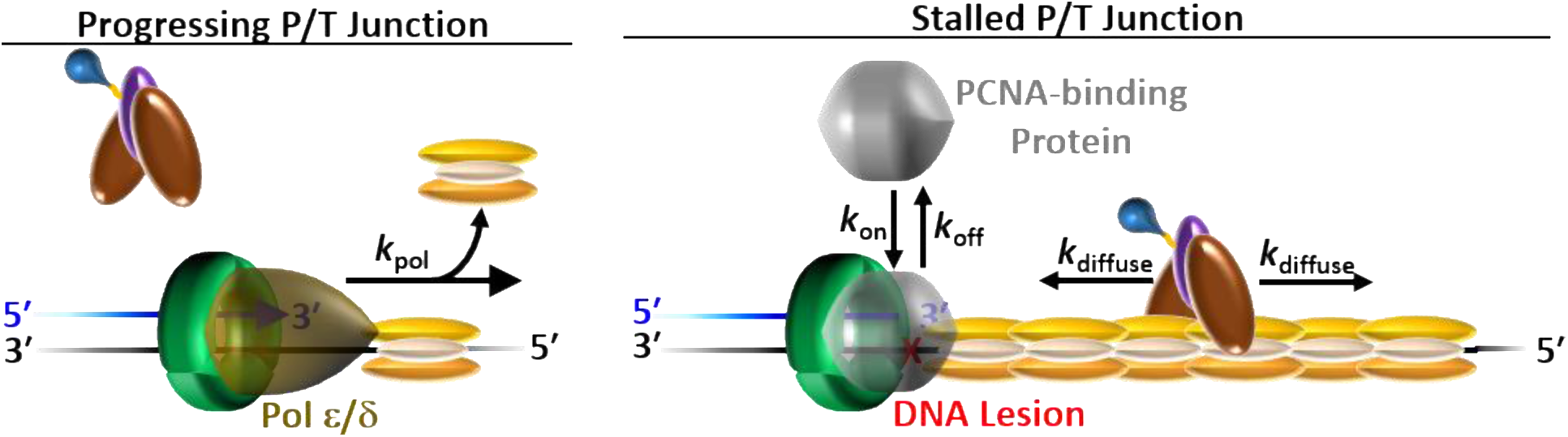
RPA interacts with Rad18 to regulate PCNA monoubiquitination. (*Left*) The short RPA filaments adjacent to progressing P/T junctions are rapidly displaced by the replicative pols ε and δ. Here, interactions of Rad18 with RPA and PCNA on ssDNA are prohibited and Rad6/Rad18 complexes remain disengaged. (***Right***). The long, persistent RPA filaments generated at P/T junctions stalled at DNA lesions promote Rad18•RPA interactions and localize a Rad6/Rad18 complex to stalled P/T junctions independently of PCNA binding. Once engaged, the Rad6/Rad18 complex translocates randomly along the RPA filament by thermal-driven diffusion. These movements selectively elevate the relative frequency of collisions between the loaded PCNA target and the Rad6/Rad18 complex. Together, these Rad18•interactions promote monoubiquitination of PCNA encircling stalled P/T junction.

In the current smFRET setup (**Fig. 2A**), dissociation of a Rad6/Rad18 complex from an RPA filament cannot be defined. Thus, it is unknown how far a Rad6/Rad18 complex can diffuse along an RPA filament before dissociating into solution. Extensive diffusion would ensure monoubiquitination of PCNA encircling a stalled P/T junction regardless of where the Rad6/Rad18 complex initially engaged the adjacent RPA filament. However, this mechanism would likely be impacted by collisions with other proteins bound to the RPA filament, resulting in local trapping of the Rad6/Rad18 complex on small segments of the RPA filament^37–40^. Thus, we envision that a Rad6/Rad18 complex diffuses along short segments of the RPA filament (up to 6 RPA molecules, **Fig. 1**) before dissociating into solution. Given the wide distribution of UVR-induced lesions after exposure to physiologically-relevant fluences (1 lesion every 3 – 4 kilobases) and the high, local concentration of RPA molecules on stalled P/T junctions, the disengaged Rad6/Rad18 complex likely re-associates with the same RPA filament at a random position. In this model, intermittent dissociation events between diffusive translocations allow a Rad6/Rad18 complex to escape local trapping and bypass proteins bound to the RPA filament. However, short diffusion lengths require that a Rad6/Rad18 complex must ultimately engage the RPA filament at a position near the stalled P/T junction in order to monoubiquitinate the resident PCNA. This proposed model is the focus of ongoing studies.

## Methods

### Oligonucleotides

DNA constructs were synthesized by Integrated DNA Technologies (Coralville, IA) and purified on denaturing polyacrylamide gels. Concentrations of unlabeled DNAs were determined from the absorbance at 260 nm using the calculated extinction coefficients. For DNA labeled with a cyanine dye, concentrations were determined from the absorbance at 550 nm (for Cy3) or 650 nm (for Cy5) using the extinction coefficient of the respective dye. Primer/Template (P/T) DNA substrates were annealed by mixing the primer and equimolar amounts of complementary templates strand in 1X Annealing Buffer (10 mM Tris-HCl, pH 8.0, 100 mM NaCl, 1 mM EDTA), heated to 95 °C for 5 min, and allowed to slowly cool to room temperature. All DNA substrates utilized in the present study are depicted in **Supplementary Fig. 1**.

### Recombinant human proteins

Wild type-PCNA, a site-specifically labeled Cy5-PCNA, Rad6, Rad6/Rad18, RFC, RPA, Ube1, and fluorescein-labeled ubiquitin (Fl-Ub) were expressed and purified as previously described^1–5^. The concentration of the purified Rad6/Rad18 complex was determined from the extinction coefficient (ε_280_ = 68570 M^−1^cm^−1^) assuming a stoichiometry of Rad6⚫(Rad18)_2_^41^ and the concentration of Rad6 within the complex was confirmed by Bradford assay using BSA (VWR) as a standard. The concentration of active RPA was confirmed as previously described (**Supplementary Fig. 2**)^17^. The plasmid (pET28-NHis-SUMO-Rad18) for the expression of Rad18 was a generous gift from Dr. Jun Huang (Life Sciences Institute, Zhejiang University, Hangzhou, China)^42^. Rad18 was expressed in *E. Coli* and purified by via slight modifications of published protocols (see **Supplementary Information**). Rad18 concentration was determined via Bradford assay using BSA as a standard.

### Protein labeling for smFRET measurements

The N-terminus of Rad6 was labeled with Cy5 (GE Healthcare). Briefly, the solution of NHS-ester functionalized Cy5 in DMSO was added dropwise under stirring conditions to a solution of Rad6 in 10 mM HEPES, pH 7.5 containing 468 mM NaCl, 2 μM ZnCl_2_ and 1 mM TCEP. The final protein:dye ratio was 1:1.1 and the labeling reaction was incubated overnight at 4° C. Labeled Rad6 was separated from free Cy5 dye by dialysis against 10 mM HEPES buffer (pH 7.5) twice. Finally, the solution was concentrated and washed twice with the storage buffer (10 mM HEPES, pH 7.5, 468 mM NaCl, 2 μM ZnCl_2_, 1 mM TCEP) via centrifugal filtration (Amicon, 3kDa MW cutoff) and stored at −80° C. The labeling efficiency was calculated by dividing the concentration of Cy5 by the concentration of Rad6. The former is determined from the absorbance at 650 nm using the extinction coefficient for Cy5. The latter is determined by Bradford assay using unlabeled Rad6 as the standard and correcting for the absorbance of Cy5 at 595 nm (ε_595_ = 140,000 + 4010 M^−1^cm^−1^). On average, each Rad6 contains one Cy5 dye (labeling efficiency = 1.10 + 0.08 Cy5/Rad6). SDS-PAGE analysis of Cy5-Rad6 indicated a single labeled species (**Supplementary Fig. 3**). Together, this indicates that Rad6 is uniformly labeled with a single Cy5 dye/protein.

### Ensemble FRET measurements

All experiments were performed at room temperature (23 ±2) °C in 1X ubiquitination buffer (25 mM HEPES, 125 mM KOAc, 10 mM Mg(OAc)2) supplemented with 1 mM TCEP. The ionic strength was adjusted to 200 mM by the addition of appropriate amounts of KOAc. First, a solution containing 110 nM of a Cy3-labeled P/T DNA substrate **(Supplementary Fig. 1**), NeutrAvidin (Thermo Scientific, 440 nM), and ATP (1 mM), was pre-incubated with excess RPA such that the concentration of free RPA is 550 nM. Cy5-PCNA (100 nM homotrimer) and RFC (100 nM) are sequentially added and retention of PCNA on DNA at equilibrium is monitored via FRET as described as previously^6,7^.

*Ubiquitination Assays.* All ubiquitination assays are performed at room temperature (23 ± 2 °C) in 1X ubiquitination assay buffer (25 mM HEPES, pH 7.5, 10 mM Mg(OAc)_2_, 125 mM KOAc) supplemented with 1 mM TCEP, and the ionic strength is adjusted to physiological (200 mM) by the addition of appropriate amounts of KOAc. All concentrations indicated below are final (i.e., after mixing). Unless indicated otherwise, experiments were performed as described previously with minor changes^10,32^. In solution A, PCNA (700 nM homotrimer) is preloaded by RFC (700 nM + 0.5 mM ATP) onto a P/T DNA substrate (700 nM + 2.8 μM NeutrAvidin) in the presence of excess RPA such that the ssDNA is saturated with RPA and the concentration of free RPA is 1.0 μM. Under these conditions, all PCNA is loaded onto the DNA and stabilized^8,9^ and, hence, the concentration of loaded PCNA is 700 nM. In solution B, Rad6/Rad18 (7 nM heterotrimer) is pre-incubated with Ube1 (14 nM + 0.5 mM ATP) and Fl-Ub (4.55 μM) for 10 minutes. Under these conditions, all Rad6/Rad18 is charged with ubiquitin^32^ and, hence, the concentration of charged Rad6/Rad18 is 7 nM. Ubiquitination of target proteins is initiated by mixing equal volumes of solutions A and B. Aliquots are removed at the indicated time points and quenched 1.33-fold into 1X reducing loading buffer (5 mM Tris, pH 6.8, 7.5% glycerol v/v, 0.375% SDS, 0.51 M β-mercaptoethanol, Bromophenol Blue). Under these reducing and denaturing conditions, only proteins containing covalent isopeptide bonds with ubiquitin are observed. After all time points are completed, samples are analyzed by fluorescence scanning as described previously to yield the concentration of monoubiquitinated PCNA clamps ([P]_T_ = [E•P] + [P]) ^10^. Data points were divided by the initial concentration of pre-charged Rad6/Rad18 ([E]_0_ = 7 nM) and plotted as function of time. For each condition, the steady state phase is fit to the equation^16^ 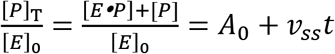. where *v_ss_* = *k_mu,obs_ k_cat_* / (*k_mu,obs_* + *k_cat_*) and *A*_0_ = [*k_mu,obs_* /(*k_mu,obs_* + *k_cat_*]^2^. Values for *k*_mu,obs_ and *k*_cat_ are calculated from the values for A_0_ and *k*_cat_ reported in **Table 1**.

### smFRET measurements

Quartz microscope slides (Finkenbeiner, USA) were thoroughly cleaned as previously described^43^ and slide surfaces were coated with polyethyleneglycol (PEG) and PEG-biotin at a 99:1 ratio. First, the BioCy3P/T-70ss DNA substrate (**Supplementary Fig. 1**) was immobilized on a microscope slide surface via biotin/streptavidin conjugation and then pre-incubated with 0.5 μM RPA for 10 minutes followed by a wash. Next, a solution containing 10 nM Cy5-Rad6, 20 nM Rad18, 1.6 mM protocatechuic acid (PCA, HWI Pherma Services), 0.16 units/mL protocatechuate-3,4-dioxygenase (PCD, Sigma) and 1 mM trolox (Sigma, MO, USA) was injected. After a 10 minute incubation to deplete oxygen, two-color smFRET measurements were performed using a prism-coupled total internal reflection fluorescence (TIRF) microscope system that is based on a Nikon TE2000 microscope (Nikon, Japan) as previously described^43^.

Briefly, the slide surface was illuminated with a 532 nm laser through a prism mounted on top of the slide. Fluorescence emission was collected through a water immersion objective lens (Nikon, Plan Apo, 60x, 1.2 NA) and bifurcated to two different paths to separately image donor (Cy3) and acceptor (Cy5) signals on an EMCCD camera (Cascade-II, Photometrics). A time-series stack of fluorescence images with 150 ms signal integration was recorded until ~70% Cy3 spots are photobleached. Several stacks of images were recorded focusing on different regions of the slide surface. The intensities of Cy3 emission and corresponding Cy5 emission were obtained from the stacks of images. The background fluorescence signal after photobleaching was taken as the zero-fluorescence level and subtracted from the fluorescence signal. From the relative intensities of Cy3 and Cy5, the FRET efficiencies were estimated with I_Cy5_/(I_Cy3_ + I_Cy5_), where I is the fluorescence intensity. From the dynamics of the FRET efficiency levels, the time windows of the FRET-on states were defined. We first identified FRET-on events with a 4-frame average of FRET efficiency ≥ 0.1 which was also verified by visual inspection. The first and last points of each event that show anti-correlated Cy3 and Cy5 intensities were defined as the starting and ending points of a τ_on_ window. We observe a total of 88 τ_on_ events over 1100 s of the 362250 s total observation time. Under these conditions the probability of observing a double-binding event is negligible (~0.0009 %).

## Acknowledgements

We would like to thank members of the Hedglin, Lee, and Benkovic labs for helpful insights, discussion, and critical reading of this manuscript. This work was supported in part by the NIH (grants RO1 GM123164 to T.H.L. and RO1 GM13307 to S.J.B.)

## Author Contributions

M.H., T.H.L. designed research. M.L. and B.S. performed research. S. J.B. contributed new reagents. M.H., T.H.L., M.L., and B.S. analyzed data. M.L., B.S., M.H., T. H.L. and S.J.B. wrote the paper.

