## Supplementary Information for "PCNA monoubiquitination is regulated by diffusion of Rad6/Rad18 complexes along RPA filaments"

### Supplementary Methods

*FRET-based active site titration of human RPA.* The concentration of active RPA was calculated via a FRET-based assay, as previously described[1]. The Cy3/Cy5-labeled ssDNA oligonucleotide (poly(dT)30-FRET) was synthesized by IDT (Coralville, IA) and is shown in **Supplementary Fig. 1**. All experiments were performed at room temperature (23 ±2 °C) in 1X ubiquitination buffer (25 mM HEPES, 125 mM KOAc, 10 mM $Mg(OAc)_2$) supplemented with 1 mM TCEP. The ionic strength was adjusted to 200 mM by the addition of appropriate amounts of KOAc. A Jobin Yvon FluoroMax-4 Fluorimeter was used to excite Cy3 and observe Cy3 and Cy5 emissions. The excitation and emission slit widths were all set to 5 nm. The Cy3 excitation wavelength was set to 514 nm and fluorescence emission intensity ($I$) was measured at 561 nm (Cy3 FRET donor fluorescence emission maximum) and 665 nm (Cy5 FRET acceptor fluorescence emission maximum). The Cy3/Cy5-labeled ssDNA (10 nM) was titrated with increasing concentrations of human RPA and the FRET value ($I_{665}/I_{561}$) at each RPA concentration was calculated by dividing the fluorescence emission intensity at 665 nm ($I_{665}$) by the fluorescence emission intensity at 561 nm ($I_{561}$)[2]. At each RPA concentration, FRET values were recorded every minute until the FRET signal maintained a constant value for at least 2 minutes and data points within this region were averaged to obtain the final FRET value.

*Rad18 expression and purification.* BL21-(DE3)-RP (Agilent Technologies) *E. coli* cells transformed with pET28-N-His-SUMO were plated on LB agar plates supplemented with Kanamycin (30 mg/ml) and incubated at 37$^{o}$C. After 12 – 15 h, 130 mL of LB media supplemented with Kanamycin (30 mg/ml) was inoculated with a single colony and incubated overnight (O/N) with aeration at 37$^{o}$C. After 12 – 15 h, each of 12, 1L LB media cultures supplemented with Kanamycin (30 mg/ml) was inoculated with 10 mL of the O/N culture and incubated at 37 $^{o}$C with aeration. When the $A_{600}$ of the 1L cultures reached ~0.60, the temperature was decreased to 15 °C, the cultures were induced by the addition of IPTG (0.25 mM) and incubated at 15$^{o}$C with aeration. After 15 – 17 h, cells were collected by centrifugation at 4$^{o}$ C. The resultant cell pellet was re-suspended in 3.6 mL of Lysis Buffer (10 mM HEPES, pH 7.5, 150 mM NaCl, 2 μM ZnCl2, 0.5 mM TCEP) per gram of cell pellet. Lysis buffer was supplemented with 1 EDTA-free complete protease inhibitor cocktail tablet (Roche) per 50 mL of lysis buffer. Cells were lysed by sonication and cell lysates were clarified by centrifugation at 4 °C. Clarification was carried out at 4°C and EDTA-free complete protease inhibitor cocktail tablets (1 per 50 mL of solution) were included in all buffers for cobalt-affinity chromatography.

The supernatant was applied to 10 mL cobalt agarose beads (high-density) (GoldBio) that were pre-equilibrated with lysis buffer and the resultant slurry was gently rocked. After 1 h, the resin was collected by centrifugation, resuspended Lysis Buffer, and applied to a column. The packed beads were washed stepwise with 5 column volumes (CV) of Lysis Buffer supplemented with; 1) 5 mM imidazole; 2) 10 mM imidazole, and; 3) 20 mM imidazole. Bound protein was then eluted with 10 CV of Lysis Buffer supplemented with 500 mM imidazole. Eluted samples were pooled and loaded at 1 mL/min onto a 5-mL Heparin column (GE Healthcare Life Science) that had been pre-equilibrated with Buffer A (10 mM HEPES, pH 7.5, 2 μM ZnCl2, 0.5 mM TCEP) supplemented with 150 mM NaCl. The column was washed stepwise with 3 CV of Buffer A supplemented with; 1) 200 mM NaCl; 2) 250 mM NaCl and; 3) 300 mM NaCl. The column was washed with 10 CV Buffer A supplemented with 350 mM NaCl and then bound protein was eluted with a 38 CV linear NaCl gradient (350 – 660 mM) in Buffer A. Fractions containing Rad18 eluted at ~ 468mM NaCl and were pooled and concentrated via centrifugal filtration (Amicon, 30 kDa MW cutoff). The concentrated protein was divided into aliquots, frozen in liquid nitrogen, and stored at –80°C. The concentration of full-length Rad18 was determined via Bradford assay using BSA as a standard.

### Supplementary Figures

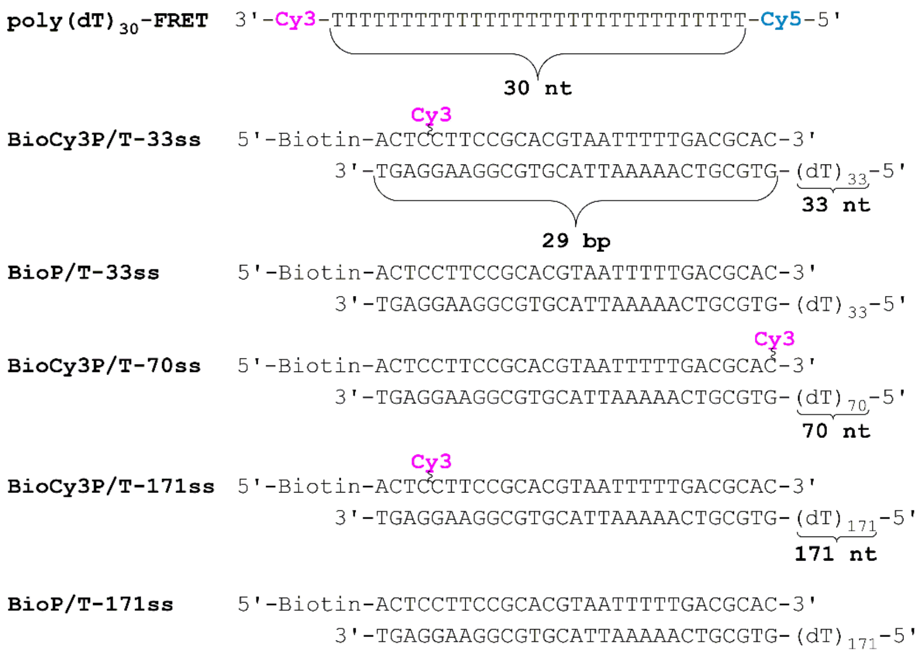


**Supplementary Fig. 1**. DNA substrates utilized in this study. The sequences and lengths of the double-strand DNA (dsDNA) regions (29 bp) of each DNA substrate are identical. All single-strand DNA (ssDNA) regions are comprised only of T (i.e., poly(dT)$_X$) and are incapable of adapting stable secondary structures[3]. When annealed, each DNA substrate mimics a blocked primer/template (P/T) junction. The size of the double-stranded P/T region (29 bp) is in agreement with the requirements for assembly of a PCNA ring onto DNA by RFC[45,6]. The single-stranded DNA (ssDNA) regions adjacent to the 3′-end of the P/T junctions accommodate 1 – 6 RPA molecules which prevents loaded PCNA from sliding off the ssDNA end of the substrate[5,6]. When pre-bound to NeutrAvidin, the biotin attached to the 5′-end of a primer strand prevents loaded PCNA from sliding off the dsDNA end of the substrate. A Cy3 dye near the 5′-end of a primer strand serves as a FRET donor.

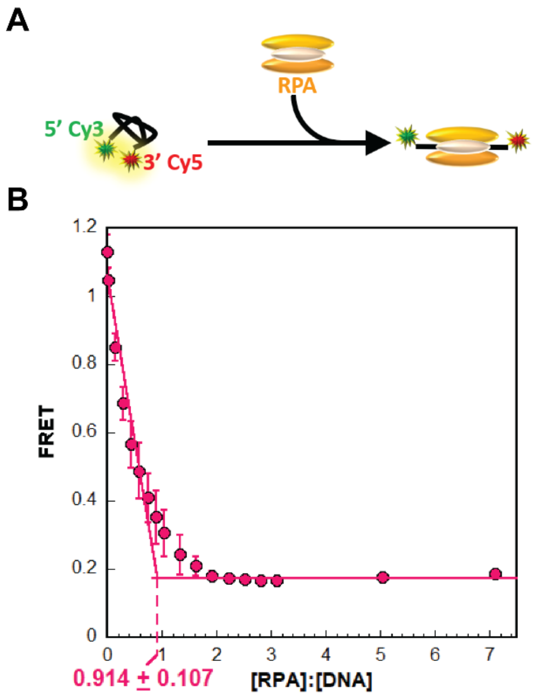


**Supplementary Fig. 2.** FRET-based active site titration of RPA. (**A**) Schematic representation of the FRET-based assay. In the absence of RPA, free DNA (poly(dT)30-FRET) forms a compact, flexible structure, bringing the two cyanine fluorophores close together and yielding a high FRET. The poly(dT)30-FRET substrate can accommodate a single RPA and binding of RPA stretches the engaged ssDNA and increases its bending 2 – 3 -fold[1,7], thereby increasing the Cy3–Cy5 distance and reducing FRET. (**B**) RPA titration. Poly(dT)$_{30}$-FRET (10 nM) is titrated with RPA and FRET is monitored. The observed FRET ($I_{665}/I_{561}$) is plotted as a function of the [RPA]:[DNA] ratios (each corrected by respective dilution factors) and each data point represents the average $\pm$ standard deviation of at least three independent measurements. Under these experimental conditions, binding is stoichiometric and, hence, FRET decreases linearly until the ssDNA is saturated with RPA (i.e., the equivalence point). Data is fit to two segment lines (a linear regression with a negative slope and a flat line) and the equivalence point (indicated) is calculated from the intersection of the two segment lines. Saturation is reached at approximately one RPA heterotrimeric complex per poly(dT)$_{30}$-FRET ssDNA (0.914 $\pm$ 0.107), indicating that RPA is fully active.

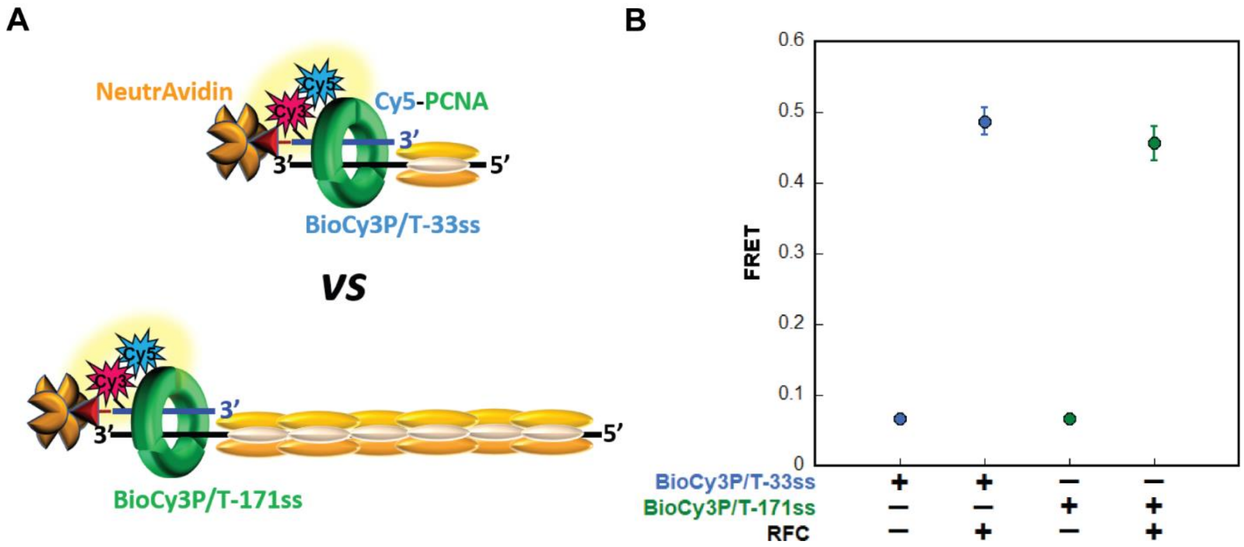


**Supplementary Fig. 3**. Retention of PCNA on DNA. (**A**) Schematic representation of the FRET experiments. Cy5-PCNA was assembled on BioCy3P/T DNA substrates in the presence of RPA, and FRET was monitored at equilibrium. (**B**) FRET in the presence of RPA. FRET is only observed when the human clamp loader, RFC, is included. FRET values measured in the presence of RFC are within experimental error for the BioCy3P/T-33ss (**Blue**, 0.487 $\pm$ 0.0194) and BioCy3P/T-171ss (**Green**, 0.456 $\pm$ 0.0246) DNA substrates, indicating that the same amount of PCNA is loaded onto and stabilized at a P/T junction in the same FRET state (i.e. orientation), regardless of the length of the adjacent RPA filament.

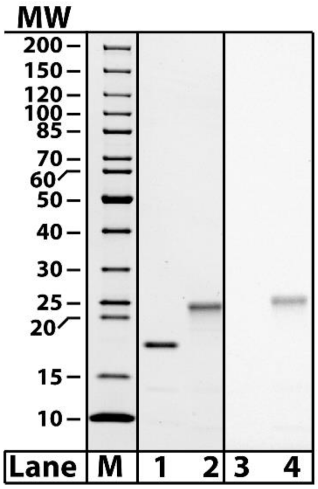


**Supplementary Fig. 4**. SDS-PAGE analysis of Cy5-labeled Rad6. Rad6 (Lanes 1 and 3, 16 pmol), and Cy5-Rad6 (Lanes 2 and 4, 16 pmol) were loaded on a SDS 4 - 20% gradient polyacrylamide gel and imaged with UV trans illumination to visualize Cy5 (lanes 3 - 4) prior to staining with Coomassie Blue (lanes 1 - 2). The MW of each band within the marker lane (M) is indicated on the left.
